# *In vitro* assays for clinical isolates of sequence type 131 *Escherichia coli* do not recapitulate *in vivo* infectivity using a murine model of urinary tract infection

**DOI:** 10.1101/2024.05.08.593128

**Authors:** Courtney P. Rudick, Rachel S. Cox, Travis J. Bourret, Nancy D. Hanson

**Author notes:** Corresponding author: (NDH).

## Abstract

Sequence Type 131 isolates are a major cause of cystitis and pyelonephritis. Many studies rely solely on *in vitro* assays to screen for bacterial virulence factors associated with the pathogenicity of clinical isolates of *E. coli*. Few studies have compared *in vitro* findings to *in vivo* infectivity of clinical isolates. The purpose of this study was to evaluate the correlation between *in vitro* assays with the ability to cause cystitis and pyelonephritis in a murine model of urinary tract infection. *In vitro* assays were conducted according to published protocols and included: motility assays, biofilm formation, epithelial cell adhesion and invasion, and curli production. Twenty-one UPEC isolates of *E. coli* ST131 and non-ST131 were used for both *in vivo* and *in vitro* studies. Six mice per isolate were inoculated via urethral catheterization. CFUs were determined from bladder and kidneys. *In vitro* and *in vivo* correlations were evaluated by multiple linear regression analysis. Pairwise linear regressions showed trendlines with weak positive correlations for motility, adhesion, and invasion and weak negative correlations for hemagglutination, biofilm and curli production. The ability of *E. coli* ST131 and non-ST131 clinical isolates to cause cystitis and pyelonephritis varies among strains. The R^2^ Pearson Correlation value was less than ±0.5 for any pair, indicating little to no statistical association between *in vitro* and *in vivo* findings. These data show *in vitro* data are not predictive of the ability of ST131 *E. coli* to infect and/or cause disease in a mouse model.

**Author summary:** Urinary tract infections affect 150 million people annually and *E. coli* ST131 have become the pandemic strain responsible for a majority of UTI, cystitis, and pyelonephritis cases. How ST131 *E. coli* have become such prolific strain still remains to be elucidated. When evaluating bacterial pathogenicity, it is customary practice to use *in vitro* assays to predict isolate virulence and mechanisms of fitness, due to the lower cost, and relative ease of experimentation compared to more costly and complicated *in vivo* models. It is also common to use model organisms like pathogenic *E. coli* CFT073 or non-pathogenic lab strains such as BW25113 as representatives for the entire species. However, our research has shown that not only are model organisms substantially different from clinical isolates of ST131 *E. coli*, but *in vitro* assays are poor predictors of clinical isolates’ ability to cause infection in a murine model of UTI. As such, research into the mechanisms of fitness for ST131 infectivity need to veer away from studying only model organisms and focus on utilizing pathogenic clinical isolates in conditions that more closely recapitulate urinary tract environmental niches.

## Introduction

Urinary tract infections (UTIs) are the second most common bacterial disease occurring in 150 million people globally every year, with women between the ages of 18-24, and women over the age of 60 at especially high risk of recurrent UTI. [1,2]. Each year, one in five women in the most at-risk age group will have a symptomatic UTI; of these infections, *E. coli* is the most common causative agent, being detected in over 75% of cases [1,3,4] *E. coli* that causes UTIs are categorized as uropathogenic *E. coli* (UPEC), and pathotypes associated with cystitis, pyelonephritis, and bacteremia are categorized as extraintestinal pathogenic *E. coli* (ExPEC) [3].

In order to colonize the urinary tract, *E. coli* must be able to evade host defenses, move from the intestinal tract to the periurethral opening, and adhere to the epithelial cells in the urethra, bladder, and kidney [5,6]. Four to twenty-four hours after bladder colonization the bacteria express adhesins including curli, type I and P-fimbriae which attach to the mannose moieties of uroplakin receptors expressed on the urethral epithelial cells which allows the *E. coli* to travel up the urethra to the bladder [7]. Cystitis occurs when ExPEC infection is localized to the bladder and is characterized by the continual expression of fimbriae by *E. coli* [7]. If fimbriae are no longer expressed, the bacteria are able to ascend up the ureters and into the kidneys inducing pyelonephritis [7]. Once in the kidneys, P-fimbriae attach to digalactoside receptors expressed on the kidney epithelial cells establishing infection [7]. Adhesins like fimbriae and curli are important in host-immune evasion and intracellular invasion of epithelial cells [8]. Most studies evaluating pathogenicity in *E. coli* have evaluated the contribution of these mechanisms using *in vitro* assays. *In vitro* assays used to evaluate pathogenicity of an isolate include motility, biofilm production, mannose binding assays, and cellular adhesion.

A major pathogenic clone in cystitis and pyelonephritis is *E. coli*, sequence Type (ST) 131. This ExPEC pandemic clone was originally discovered on three continents (Asia, Europe, and North America) in 2008, [4,6,9]. ST131 strains are multidrug resistant and harbor chromosomal and plasmid-encoded resistance mechanisms to common antibiotics used to treat UTIs including fluoroquinolones and β-lactams. The majority of data in the literature evaluate ST131 *E. coli* using whole genome sequencing, identifying genes involved in resistance and/or virulence. These types of data although useful are insufficient to predict the pathogenicity of *E. coli* ST131. Animal studies have been performed using ST131 *E. coli* to evaluate their pathogenicity but the route of infection was not through the urinary tract [10–12]. In addition, studies have used *in vitro* assays to help determine the potential pathogenicity of a particular strain [13,14]. We hypothesized that *in vitro* evaluation of the mechanisms associated with pathogenicity may not predict the ability of isolates to infect and cause disease using a mouse UTI model. To test our hypothesis, we characterized 21 clinical UPEC isolates of *E. coli* including 15 ST131, and 6 non-ST131 in both *in vitro* and *in vivo* systems to determine the correlation between these two types of analyses. Unlike studies that evaluate pathogenesis/fitness using either *in vitro* or *in vivo* methods we evaluated the same isolates using both experimental approaches. Surprisingly, little or no correlation was observed among the isolates that were successful in causing a bladder and/or kidney infection with the *in vitro* assays used to measure mechanisms associated with infectivity and pathogenicity.

## Results

### Strains used in this study

The characteristics of the strains used in this study are provided in Table 1. In addition to the sequence type, phylotype, geographical location of collection, the β-lactamase produced by the strains, and the susceptibilities to a range of antibiotics are listed. In order to compare the isolates used in this study we first determined if there were any fitness differences among the strains. Growth curves were performed and no difference in the time required to reach mid-log growth among the isolates was observed (S1 Fig). Strain CFT073 (ST73) is an isolate collected from a UTI of a patient and has been used repeatedly in the literature as a control for *in vitro* assays [14]. Knockout strains BWΔ*ompC* and BWΔ*ompF* were acquired from the Keio Collection and the double knockout clone was generously distributed by Dr Heike Brötz-Oesterhelt, University of Tübingen, Tübingen, Germany [15].

**Table 1.**
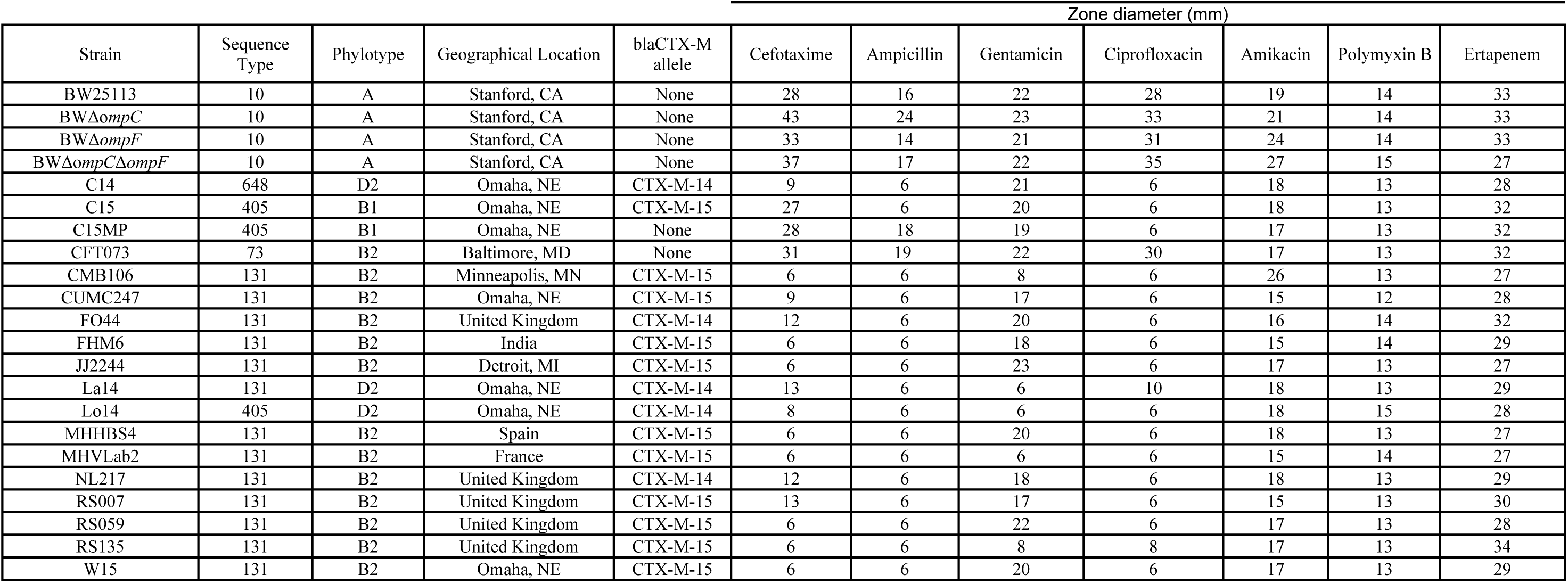
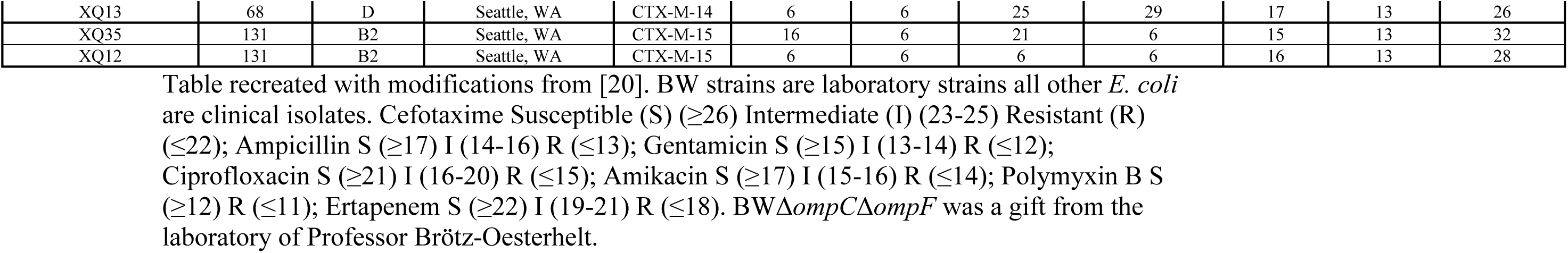
Properties of *E. coli* isolates used in this study.

### Variable pathogenicity observed among ST131 *E. coli* in a UTI mouse model

A UTI mouse model was used to establish the infectivity and pathogenicity of different strains of *E. coli* ST131 compared to non-ST131 *E. coli*. Six mice were inoculated with each *E. coli* strain using an ascending urinary tract infection model. Urine was collected from the mice up to day 28 and mice were sacrificed, and bladder and kidneys removed and evaluated. CFUs were determined for the urine, and bladder and kidney tissues. All the mice survived infection except for one. Mouse #2 in the W15 group died from peritoneal sepsis 36 hours after inoculation.

To determine the bacterial load and ability of the isolates to cause a sustained UTI, urine was collected at several timepoints throughout the study before and after inoculation. At Day 28, isolates XQ13, Lo14, and C15MP were each detectable in a single mouse per group, while C14 and most ST131 isolates (excluding CUMC247 and RS135) were detectable in high levels (up to 10^12^) in multiple mice at the end of the study (Fig 1). No lab strains were detected in urine at the end of the study. Of note, XQ12 and W15 were detectable in all mice in their respective groups throughout the course of the study (Fig 1 and S2 Fig).

**Fig 1.**
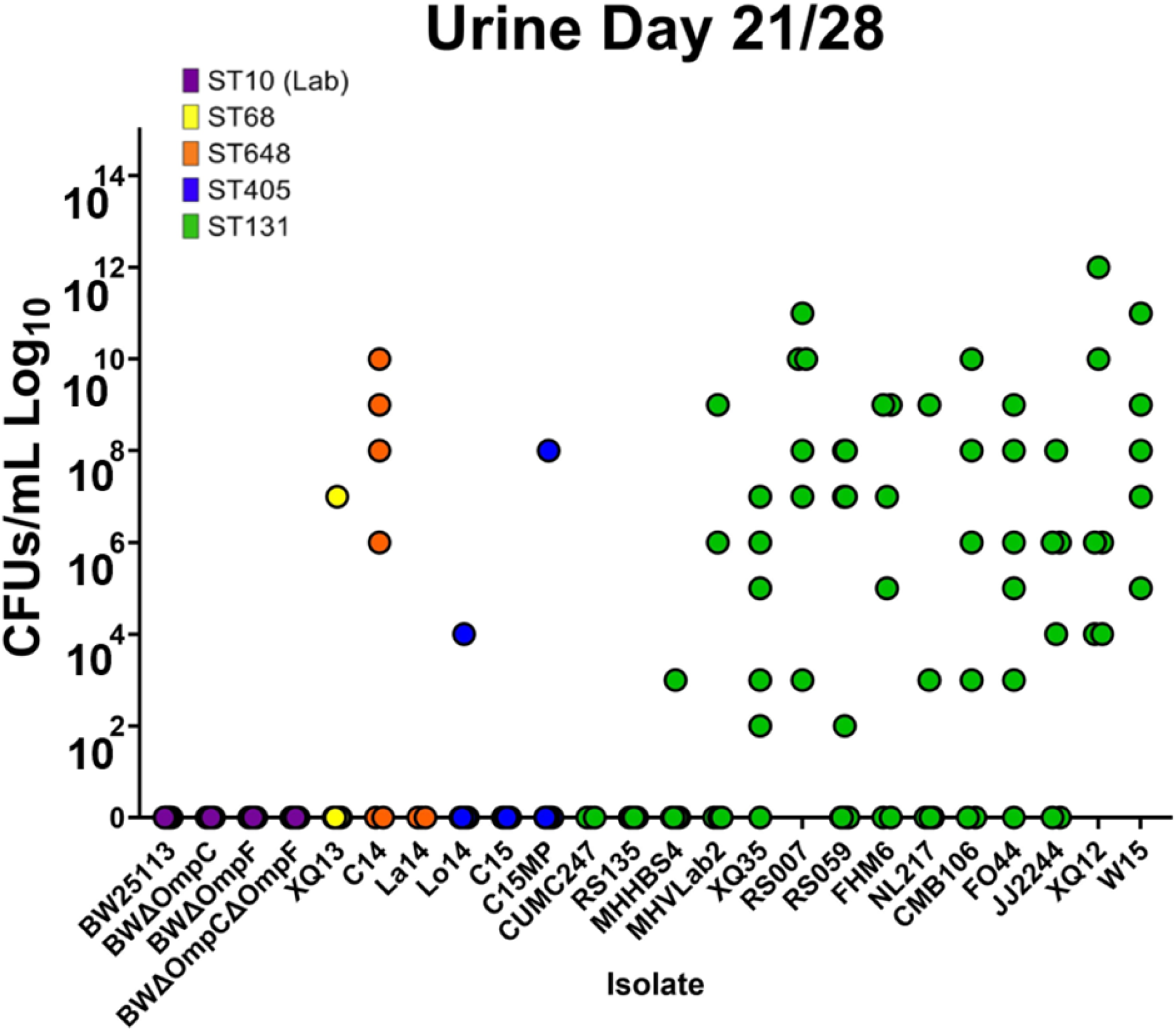
Urine colonization per mouse. CFUs from the urine of each mouse in the study on day 28, the final day of collection. Isolates XQ13, Lo14, and C15MP were all detectable in one mouse, while C14 and most ST131 isolates (excluding CUMC247 and RS135) were detectable in high levels (up to 10^12^) in multiple mice at the end of the study. Of note, XQ12 and W15 were detectable in all mice in their respective groups throughout the course of the study. Data point color represents isolate sequence type as noted in the figure legend.

The CFUs for each isolate throughout the duration of the study are shown in S2 Fig. In general, ST131 isolates caused more sustained infections with higher CFUs. Specific representative isolates have been highlighted in panels S2 Fig B-E. S2 Fig B shows, for example, infection with BWΔ*ompF* (ST10 Lab Strain) peaked at Day 3 and was rapidly cleared. W15 (ST131) started with high CFUs and increased over the course of the study (S2 Fig C). C15 (ST405) had detectable CFUs which dropped off, returned at Day 10 and Day 14, and then was cleared completely (S2 Fig D). Lastly, the burden of C14 (ST648) increased steadily until Day 10, then leveled off until the end of the study (S2 Fig E).

At 28 days, mice were euthanized with bladders and kidneys collected, weighed, homogenized, serially diluted, and plated to determine presence of infection and bacterial load per gram of the organ. The colored data points in Fig 2 represent the *E. coli* inoculum bacterial load of each mouse bladder at Day 28. None of the ST10 lab strains caused bladder infection at the end of the study. Only two non-ST131s were present in bladder tissue, C14 (ST648) and C15MP (ST405). C15MP had one positive mouse bladder with detectable, but low levels of inoculum. Most ST131 isolates were able to colonize bladder tissues at 10^3-8^ CFUs; however, ST131 strains CUMC247, RS135, and MHHBS4 were not detected in the bladder on the last day of the study. The most efficient ST131 for bladder infection was strain W15 which infected 100% of the mouse bladders with CFUs of 1 x 10^8^.

**Fig 2.**
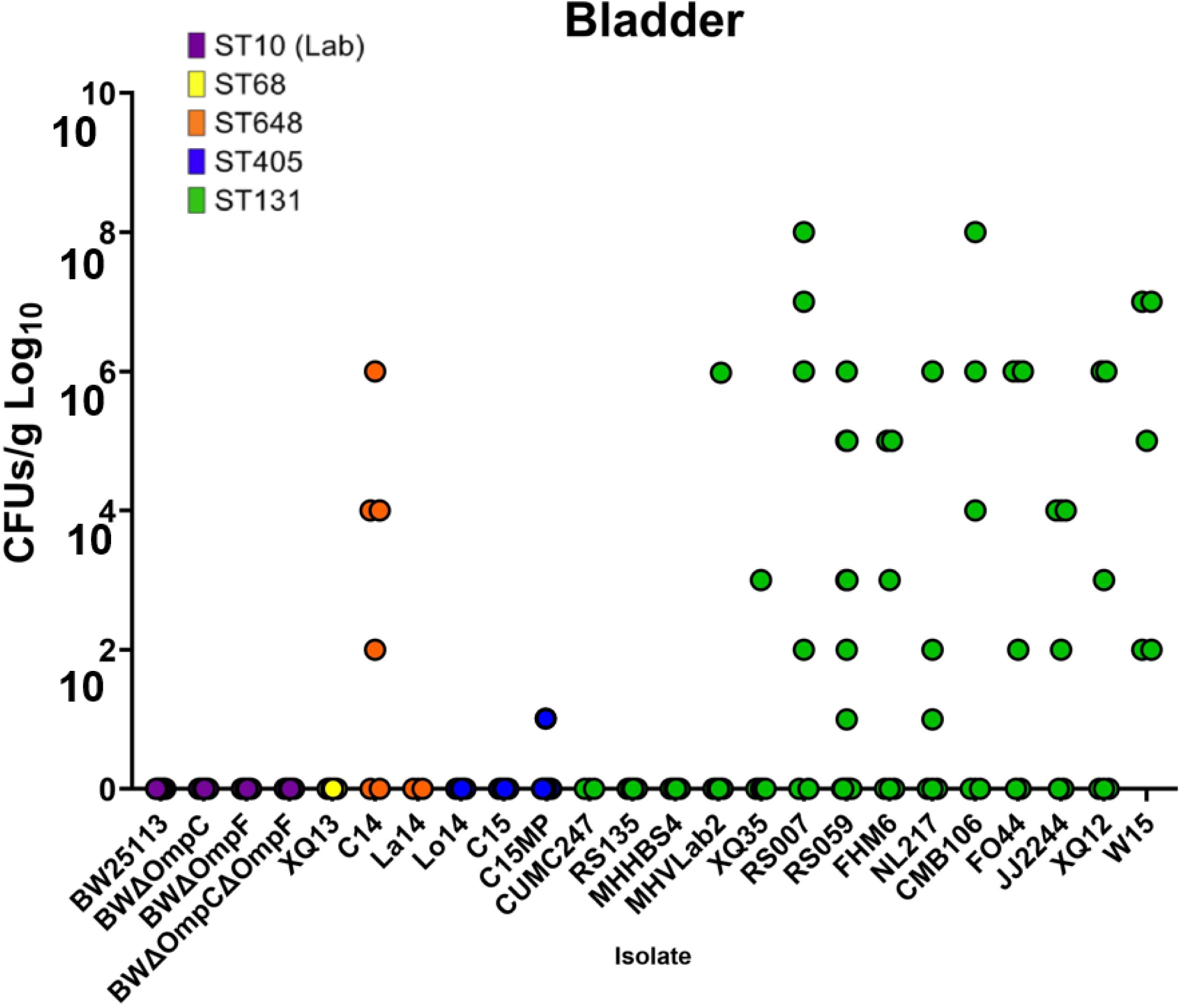
Bladder colonization per mouse. Data points represent individual mouse bladders for each strain at the level of detectable CFUs. You can see the single C15MP positive mouse bladder with detectable, but low levels of inoculum. C14 has similar infectivity as the majority of ST131 isolates in both number of bladders infected and CFUs. Three ST131 isolates (CUMC247, RS135, and MHHBS4), one ST648 (La14), and all lab strains, ST68, and ST405 strains were not detectable in any mouse bladders. Again, W15 was detectable in all mice bladder tissues. Data point color represents isolate sequence type as noted in the figure legend.

Individual mouse kidneys infected by the various strains are shown in Fig 3. The colored data points in Fig 3 represent the *E. coli* inoculum bacterial load of each mouse kidney at Day 28. In addition to infections by C14 and C15MP, CFUs were present in the kidney for non-ST131 isolates XQ13 (ST68), Lo14 (ST405) and C15 (ST405). CFUs were also detected in the kidney for ST131 isolate MHHBS4. Interestingly, in strains where the number of colonized kidneys was low, the detectable CFUs are comparable between all infected tissues. Two mice in the RS135 group showed visible changes to kidney tissues with enlarged, clear fluid filled, kidneys but no bacteria were detected from plating of tissue homogenates. As observed in the bladder, strain W15 infected the greatest number of kidneys (9/12) with CFUs of 1×10^9^.

**Fig 3.**
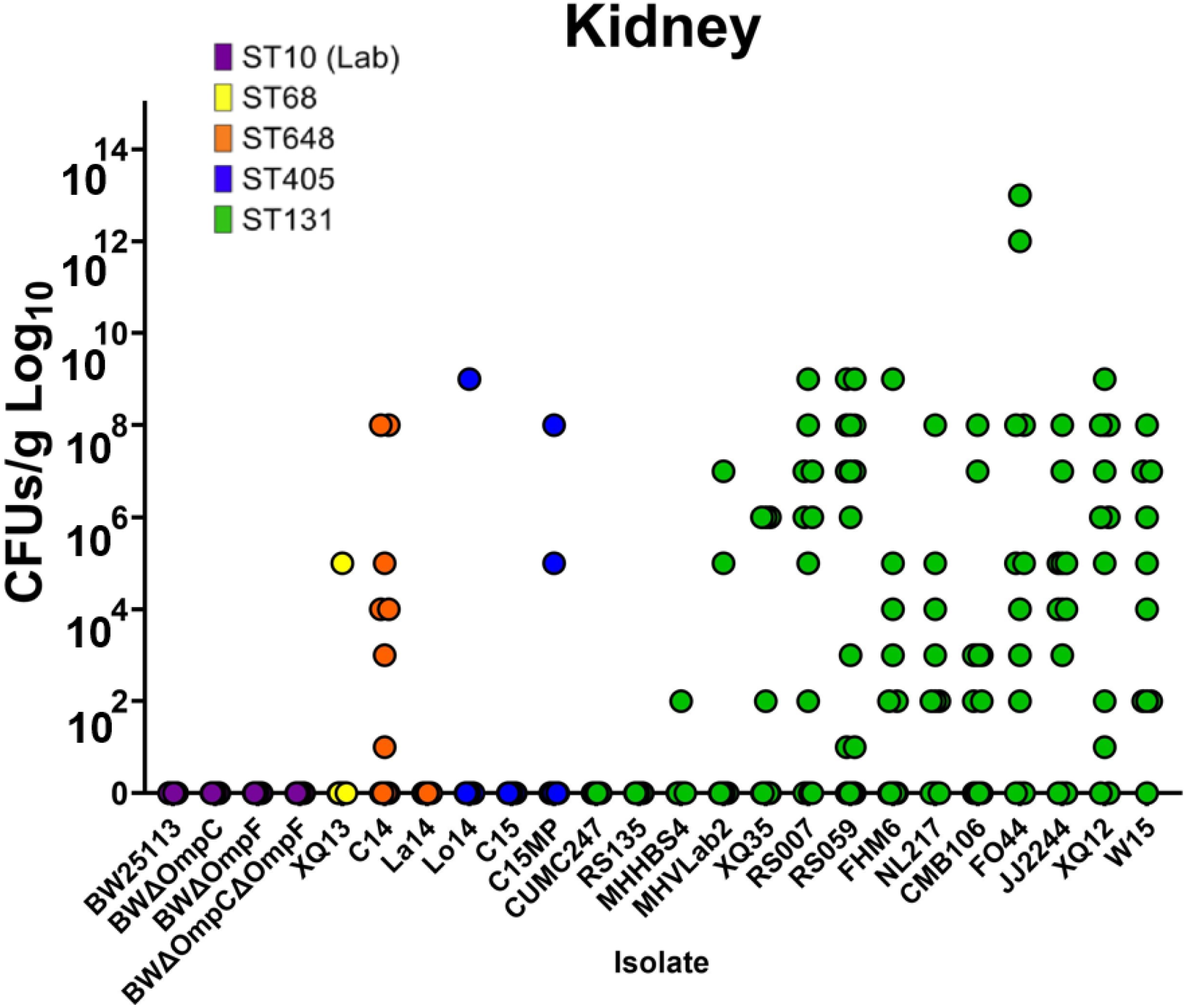
Kidney colonization per mouse. Individual points represent each mouse kidney. Almost all ST131 isolates cause infection in kidney tissues (up to 10^13^ CFUs) with C14 also infecting multiple kidney tissues at CFUs up to 10^8^. ST131 isolates CUMC247 and RS135 did not infect kidney tissues, while MHHBS4 was detectable in kidney tissues even though it was not found in bladder tissues. W15 was once again the most infective strain being found in 9/10 kidneys. Data point color represents isolate sequence type as noted in the figure legend.

### In vitro assessment of common mechanisms of fitness and virulence

*In vitro* assays used to evaluate pathogenicity of the 25 isolates included motility, biofilm production, mannose binding assays, and cellular adhesion and invasion. For all the evaluations except motility, both static and shaking cultures were used. Although these assays are typically done using static cultures, we reasoned that in a urinary tract infection it would be unlikely that the *E. coli* would exhibit a static phenotype.

#### Motility assays

Motility as determined by zone diameter of migration is depicted in Fig 4 and S3 Fig. All of the strains tested were capable of movement but to varying degrees. The non-ST-131 strains (ST68, ST648, and ST405) showed the least amount of motility among the strains tested and was equivalent to the laboratory strain, BW25113. Larger zone sizes were observed for most of the ST131 isolates indicating the ability to move. Only one ST131 strain (RS135) showed a decrease in motility compared to the other ST131 isolates and was more closely related to the movement of non-131 strains. This isolate was also incapable of infecting the bladder and the kidney using the mouse model. The lab strains deficient in porin production of either OmpC (BWΔ*ompC*) or OmpF (BWΔ*ompF*) or the double knockout strain (BWΔ*ompC* Δ*ompF*) showed similar motility to the ST131 strains. Interestingly, the OmpF knockout *E. coli*, BWΔ*ompF*, had the largest zone size at 51mm. All three *E. coli* knockout mutants showed increased motility compared to the parent strain BW25113 (<10mm). Of interest, the motility of the control strain CFT073, was similar to the non-131 strains and the laboratory strain BW but was not as motile as the ST131 strains. Although W15 was the most infective in the murine model, it only demonstrated average motility with XQ12, FHM6, NL217, and JJ2244 all being more motile while less infective.

**Fig 4.**
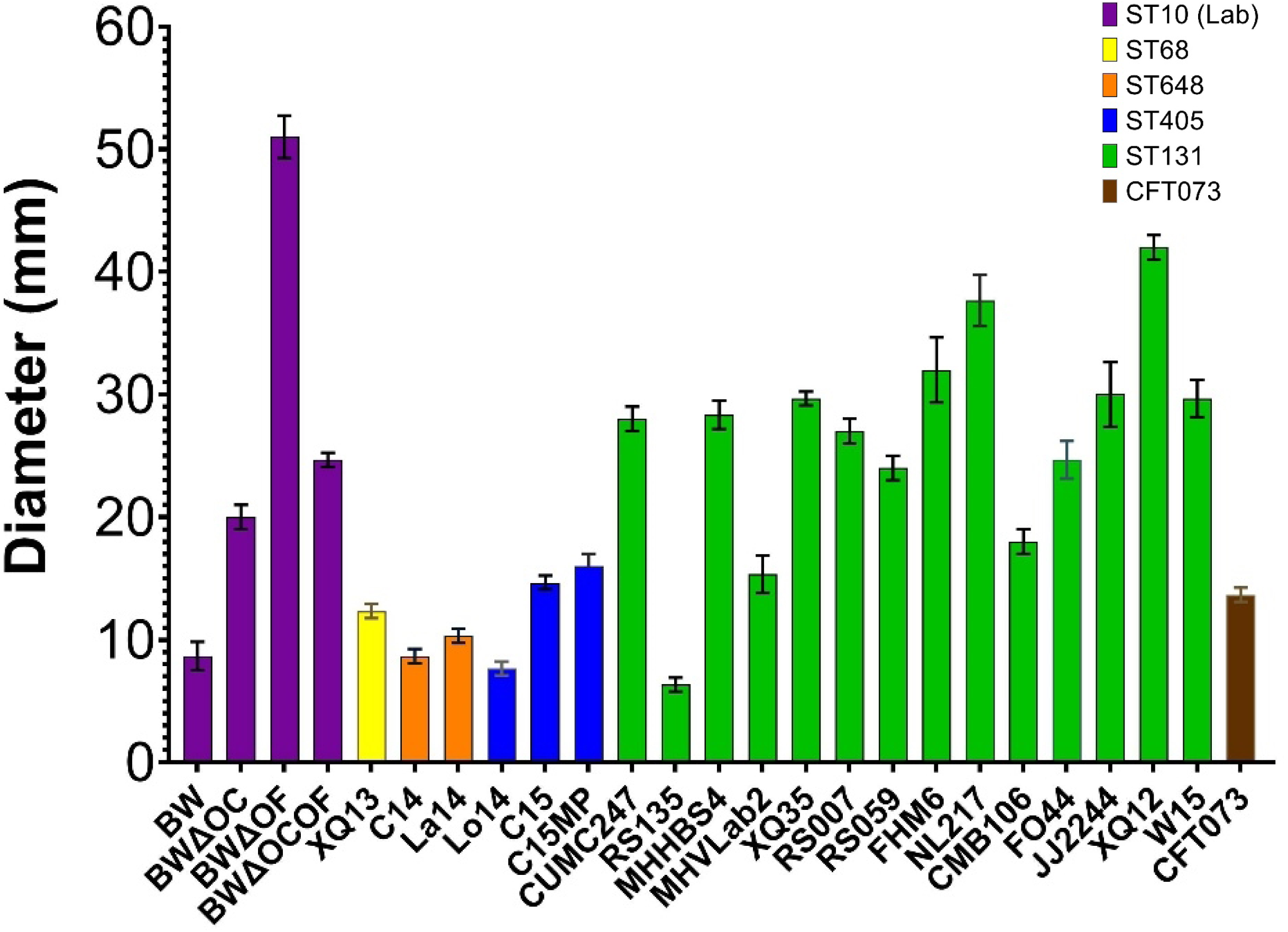
Motility of *E. coli* isolates. Zone of growth (measured across the diameter in mm) shows the ability of the isolate to move through semi-solid Agar (0.3% bactoagar in LB). Bars are colored according to isolate sequence type as defined in the figure legend. Error bars are standard deviations. Images of individual plates can be found in S3 Fig.

#### Biofilm formation

Biofilm formation is often associated with antibiotic resistance and can be considered a mechanism that contributes to pathogenicity. Although biofilm production is typically performed using static culture conditions, we evaluated biofilm production using both static or shaking conditions [16]. In general, less biofilm was produced in all isolates under static conditions (Fig 5A) as compared to shaking conditions (Fig 5B). Isolates were determined to be strong (OD>2×ODc), moderate (1.5×ODc<OD≤2×ODc), weak (ODc<OD≤1.5×ODc), or absent (OD≤ODc) as compared to the OD of a media-only well. Under static conditions most isolates were weak biofilm producers, with only MHVLab2, Lo14, W15, FHM6, and BWΔ*ompC* being moderate producers of biofilm, and only BW25113 and BWΔ*ompF* had strong biofilm production (Fig 5C). More biofilm production was noted when the organisms were grown first under shaking conditions. The exception for no change in biofilm formation between the two conditions were strains CUMC247, La14, C14, and RS135 (Fig 5C). In both static and shaking conditions biofilm production was highest for the ST10 lab strains. The double knockout mutant produced the least amount of biofilm compared to the single knockout BWΔ*ompC*. Interestingly, the BWΔ*ompF* and the parent strain BW25113 produced the most biofilm of any strain tested. The laboratory strains produced approximately 4-fold more biofilm formation in shaking cultures compared to static cultures. Interestingly MHVLab2 produced more biofilm than most ST131 isolates but demonstrated only low infectivity in either mouse bladder or kidney tissues.

**Fig 5.**
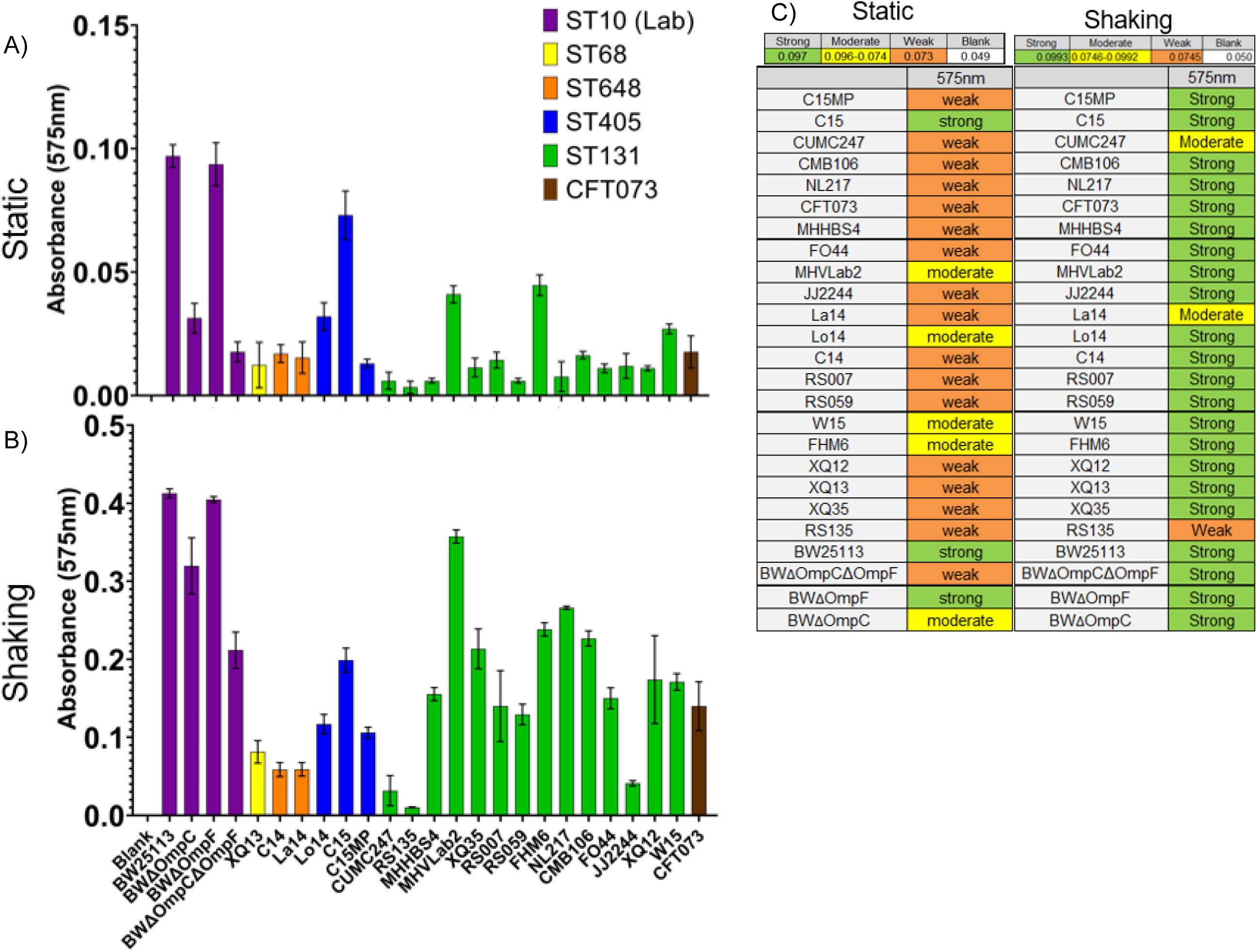
Biofilm production. Biofilm production was measured by crystal violet staining and O.D. Experiments were conducted under static (A) and shaking (B) conditions. Isolate sequence type is noted by bar color and error bars represent standard deviations. The degree of biofilm production was classified according to the following criteria: The degree of biofilm production was classified according to the following criteria: Strong (OD>2×ODc), Moderate (1.5×ODc<OD≤2×ODc), Weak (ODc<OD≤1.5×ODc), or Absent (OD≤ODc) according to [38]. Values for each classification are listed at the top of the chart in (C).

#### Cell adhesion and invasion assays

The ability of an *E. coli* cell to attach to bladder cells is one hallmark of pathogenicity. Therefore, we evaluated the ability of the *E. coli* isolates to attach or invade T24 human bladder epithelial cells when *E. coli* were grown either under static conditions or grown to mid-log immediately prior to co-culture. Laboratory BW Strains were unable to adhere to epithelial cells under either static or exponential growth conditions (Fig 6 A and B). Isolates C15MP (ST405, 5.5%) and FHM6 (ST131, 12.7%) showed the highest percentage of adherence in static conditions. When grown to mid-log phase prior to coculture, several, but not all, clinical isolates exhibited 1.5 to 2-fold higher levels of adhesion to epithelial cells. La14 (ST648) and FHM6 (ST131) showed the most increase in adhesion under mid-log conditions (1.2% to 12.6% and 12.7% to 17.3%, respectively), while C15MP (ST405), RS059 (ST131), and CFT073 (ST73) showed 2-to-4-fold lower adhesion abilities when grown to mid-log phase before co-culture. FHM6 showed the highest levels of adhesion, followed by C15MP in static conditions and La14 in Mid-log cultures, despite the inability of either C15MP or La14 to cause tissue infections in the mouse model while FHM6 effectively colonized both types of tissue.

**Fig 6.**
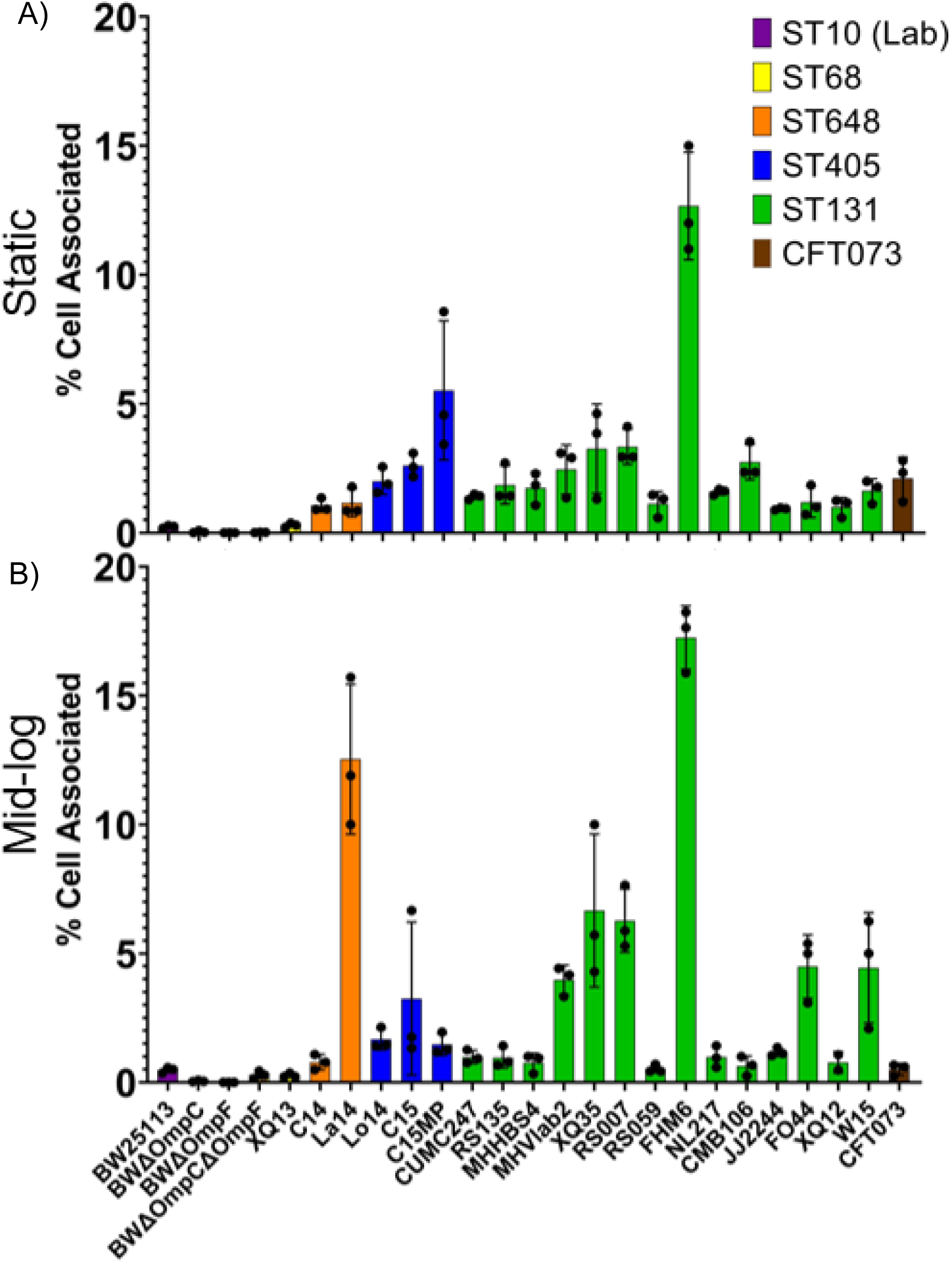
Adherence to T24 human epithelial cells. Epithelial cell adhesion was measured by determining CFUs adherent to epithelial cells as a percentage of inoculum in static (A) and shaking (B) conditions. Isolate sequence type is noted by bar color and error bars represent standard deviations.

To determine cellular invasion, the same protocol for adhesion was followed with an additional incubation step at the end adding Polymyxin B to the culture media to kill off any bacteria adhered to the outside of the cell. Fig 7 shows a smaller percentage of bacteria were able to invade cells compared to the adherence seen in Fig 6. Laboratory BW Strains were unable to invade (Fig 7 A and B) epithelial cells under either static or exponential growth conditions. Cells grown statically showed more invasion overall than cells grown to mid-log phase with ST131 isolates showing an average of 9-fold more invasion under both conditions compared to non-ST131 isolates. ST131 isolates CMB106 (0.15%) and NL217 (0.07%) showed the highest percentage of invasion in static conditions, followed by FO44 (0.03%) and XQ35 (0.02%). When grown to mid-log, most *E. coli* clinical isolates exhibited reduced or no invasion into epithelial cells, especially CMB106 (0.01%) and NL217 (0.01%). Lo14 (ST405) showed increased invasion when grown to mid-log phase (0.005% to 0.02%), and FO44 showed similar invasion at 0.03% for both growth conditions. CMB106 and NL217 demonstrated the highest percent of epithelial cell invasion (0.15% and 0.06% respectively) but were only average compared to all tested ST131s for bladder (50% and 50% respectively) or kidney (67% and 53% respectively) colonization in the mouse model.

**Fig 7.**
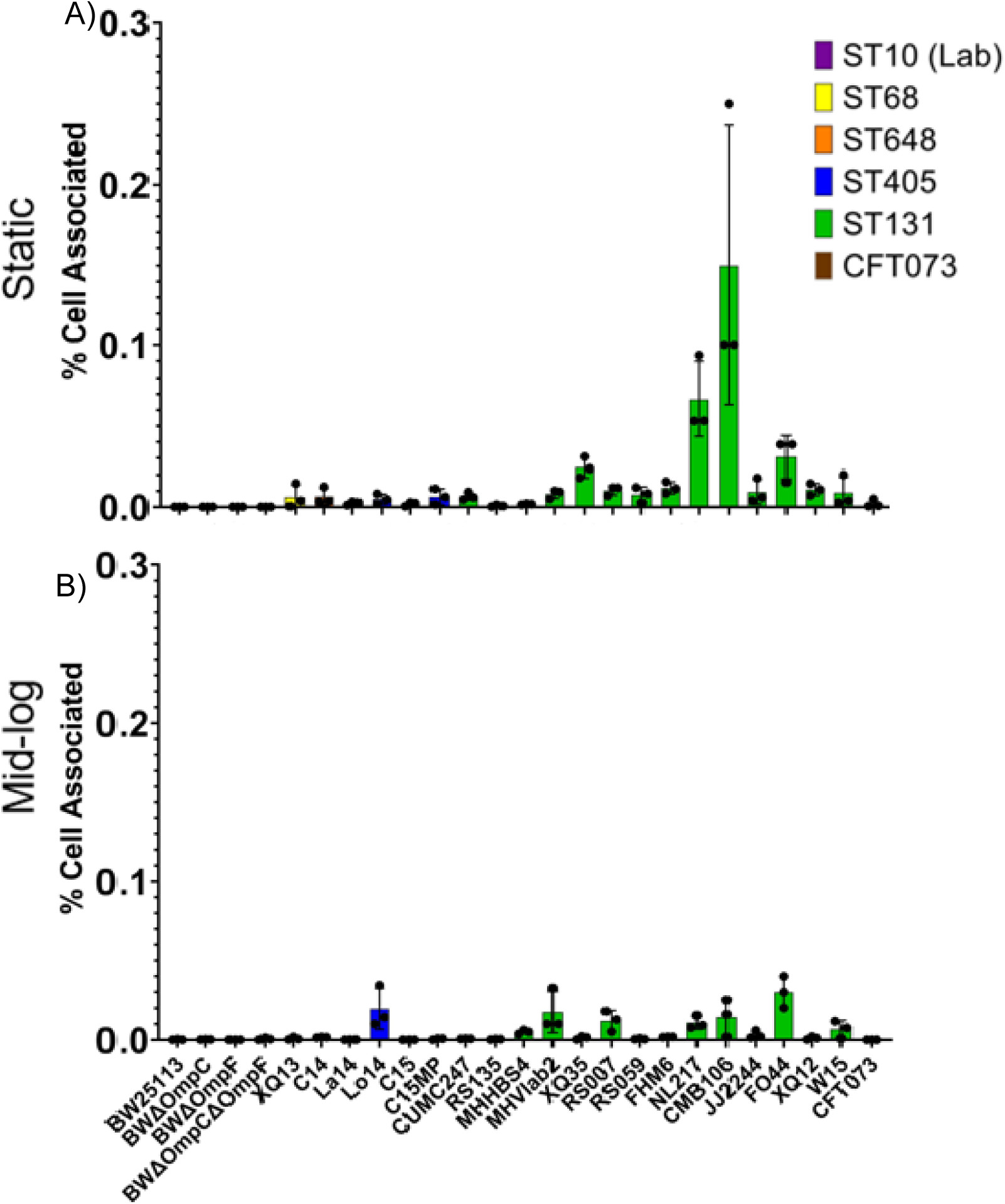
Invasion of T24 human epithelial cells. Percent of bacteria that invaded T4 bladder epithelial cells was measured by determining CFUs of bacteria present in epithelial cell cultures after incubation with Polymyxin B which kill adherent bacteria. Bacteria that invaded epithelial cells are recorded as a percentage of inoculum in static (A) and shaking (B) conditions. Isolate sequence type is noted by bar color and error bars represent standard deviations.

#### Hemagglutination assays

Type 1 fimbriae have been implicated as a pathogenic mechanism for UPEC isolates due to the FimH binding affinity for uroplakin mannosylated-glycoproteins expressed on urinary tract epithelial cells [17]. To determine the ability of the isolates to hemagglutinate red blood cells indicating the use of type 1 fimbriae, mannose sensitive hemagglutination assays were performed with bacteria grown in both static and mid-log conditions. Under static growth conditions (Fig 8A) all lab strains and isolates ST648, ST405, CFT073, and W15 exhibited mannose-sensitive hemagglutination. Interestingly, W15, the most infective isolate in the mouse model, was the only ST131 isolate capable of hemagglutination. When isolates were grown to mid-log phase no isolates were capable of hemagglutination including CFT073.

**Fig 8.**
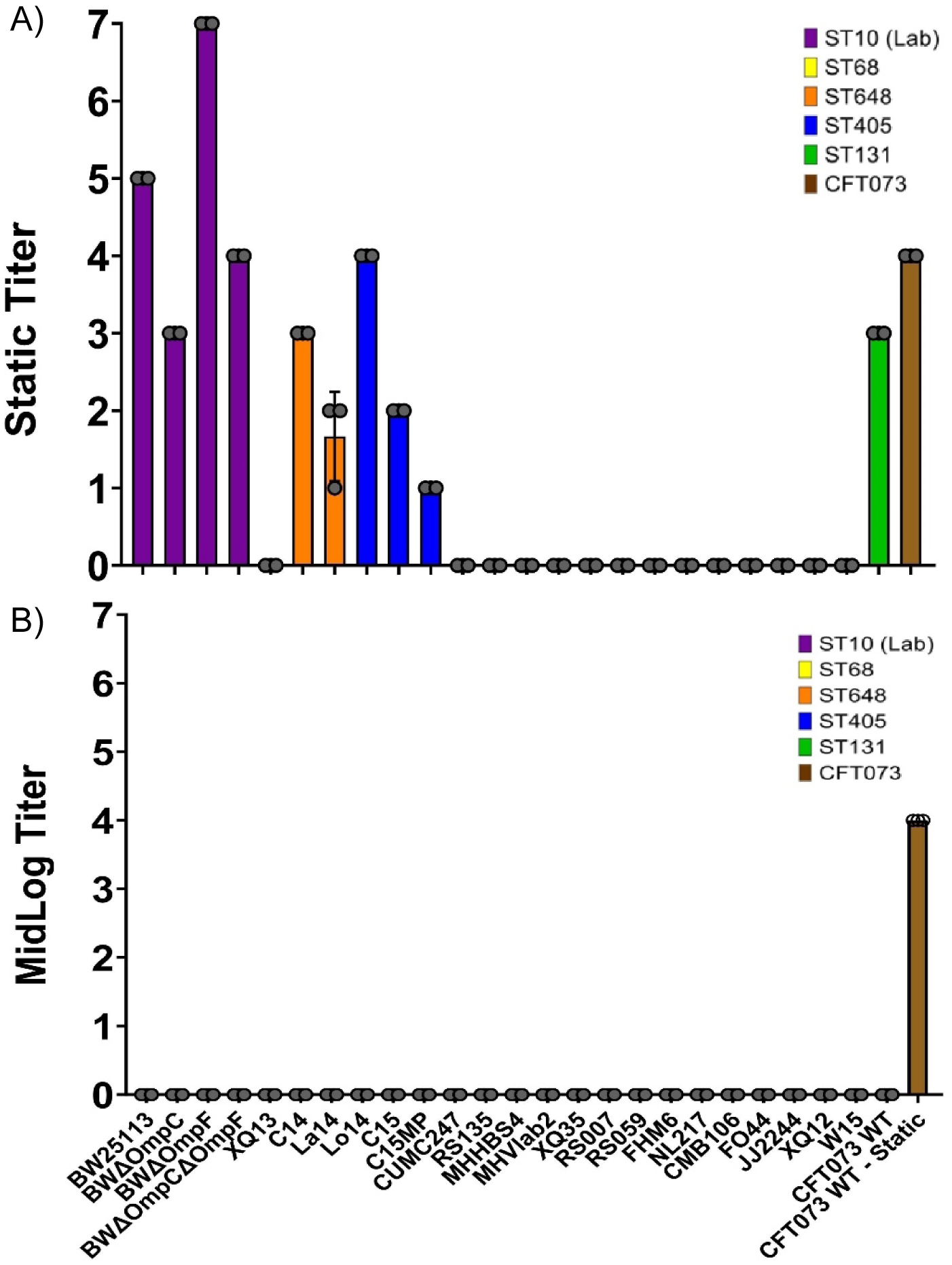
Mannose sensitive hemagglutination. Hemagglutination was measured under static (A) and shaking (B) conditions. Isolate sequence type is noted by bar color and error bars represent standard deviations.

#### Curli production

The fimbrial adhesin, Curli, encoded by the *csgA* gene is considered a virulence mechanism for uropathogenic *E. coli* due to its role in biofilm formation and cellular adhesion [18]. Congo Red assays were used to determine the level of curli production for each isolate evaluated [18]. Colonies that remained white produce no curli (Score 0-1), while colonies that produce a light color with a matte surface have some curli production (Score 2-3) and isolates with dark colored colonies and matte or wrinkled surfaces are considered high curli producers (Score 4) [18]. Both static and shaking cultures at two different temperatures (37°C and 27°C) were evaluated for curli production. Data shown in Fig 9 identified the lab strain, K12, as a high producer of Curli. As the BW25133 Lab Strains are K12 derivatives they showed similar levels of curli production to K12, which was attenuated in the OmpC knockout strain. Curli production was variable for clinical isolates of *E. coli* which as a group showed less curli production for both static and shaking cultures than lab strains at 27°C. For all clinical isolates, there was more curli production at 37°C than 27°C regardless of static or stationary growth condition. It was interesting to note that isolate, RS135, only produced curli when incubated at 37°C in either shaking or static growth conditions. XQ13 produced the most curli of all clinical isolates, despite only infecting one kidney throughout the course of the mouse study; while XQ12 and W15 showed lower levels of curli production comparable to other ST131 isolates, these isolates demonstrated higher levels of both bladder and kidney colonization in the murine model.

**Fig 9.**
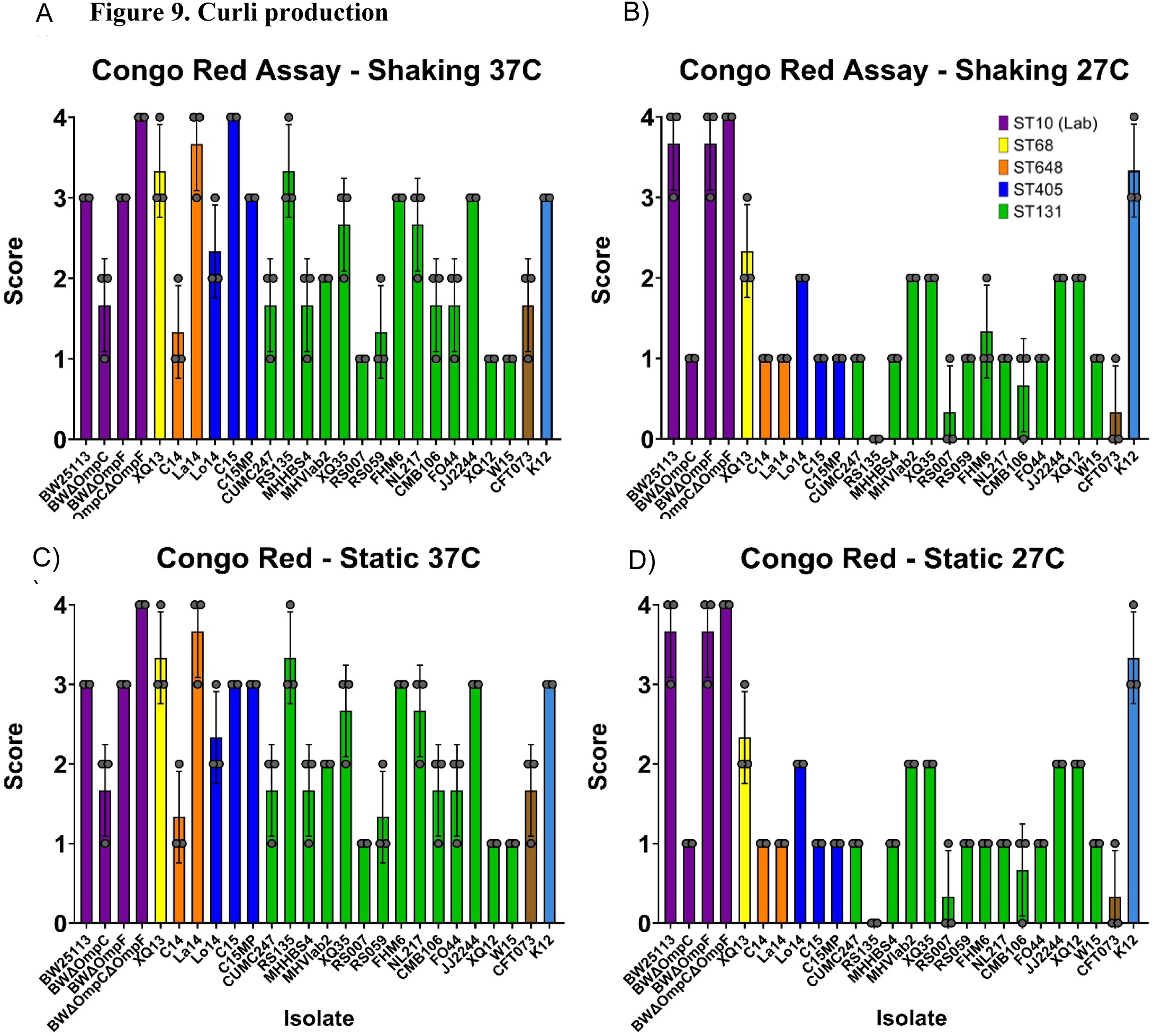
Curli production. Isolates were assessed for curli production in static (A and B) or shaking (C and D) conditions for either 24 hours at 37°C (A and C) or 48 hours at 27°C (B and D). Score 0 = an all-white colony, Score 1 = light color, Score 2 = some dark color or ring structures with a matte surface just where the color is, Score 3 = a dark, even color accompanied by a matte/dry surface, Score 4 = dark color and a rough, wrinkled colony. Isolate sequence type is noted by bar color and error bars represent standard deviations.

### Correlations between *in vitro* mechanisms of pathogenesis and *in vivo* infectivity

Phenotypic *in vitro* assays are thought to recapitulate the infectivity of isolates *in vivo*. To assess the correlation between the *in vitro* and *in vivo* data, we conducted pairwise linear regressions for each assay compared to urine cultures, bladder, and kidney infections (Fig 10 and S4 Fig). S4 Fig shows the correlations for the motility assay compared to the CFUs found in urine (S4 Fig A), bladder (S4 Fig B) and kidney during infection (S4 Fig C). While the trendline shows a weak positive correlation, the R^2^ value for the Pearson Correlation is less than 0.14 for any pair showing little to no statistical association. The correlations for hemagglutination compared to urine (S4 Fig D), bladder (S4 Fig E) and kidney (S4 Fig F) show a weak negative correlation. However, the R^2^ value for the Pearson Correlation was less than 0.15 for any pair showing little to no statistical association. The correlations for epithelial adhesion compared to urine (S4 Fig J), bladder (S4 Fig K) and kidney (S4 Fig L) show a weak positive correlation, similar to epithelial invasion (S4 Figs M-O). However, the R^2^ value for the Pearson Correlation is less than 0.1 for any pair showing little to no statistical association. While the sample variance was higher for the curli assay, the Pearson Correlation values were the strongest of any phenotypic assay with a negative correlation between curli production and kidney infection (0.2477) (S4 Fig P), bladder (0.2425) (S4 Fig Q), and urine (0.4162) (S4 Fig R). Fig 10 shows a heatmap corresponding to the strength of the Pearson Correlations for the *in vitro* assays compared to the *in vivo* mouse infection results. These results show no statistically significant relationships between any comparisons.

**Fig 10.**
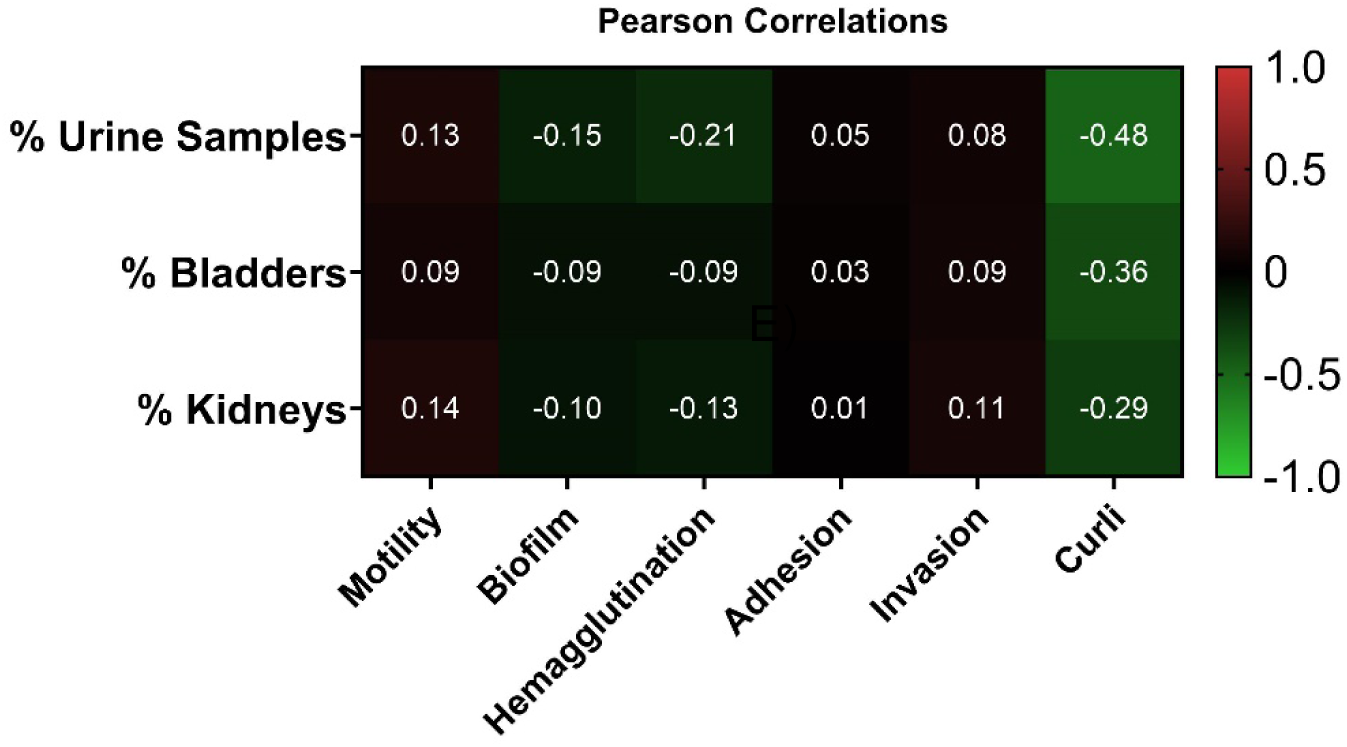
Pearson correlations for phenotypic assays and *in vivo* infection results. A heatmap of the Pearson correlations for all independent linear regressions (S4 Fig) performed comparing all *in vitro* assays to *in vivo* infectivity.

## Discussion

Mouse models of urinary tract infection are an established way to evaluate a host immune response and bacterial pathogenicity to aid in studies to improve human health [19]. However, animal studies come with several drawbacks including time, cost, and animal lives. To gain an understanding of bacterial fitness and pathogenicity, without the drawback of *in vivo* experiments, many researchers turn to *in vitro* assessments. These types of data have also been used to design diagnostic screening tools for clinical labs and healthcare settings [14]. For uropathogenic *E. coli*, *in vitro* experiments commonly include motility assays, biofilm production, curli production, adhesion/invasion assays, hemagglutination assays, and detection of bacterial enzymes like β-lactamases [14,20]. How well these experiments predict the virulence of bacteria compared to a mouse model for UTIs has not been determined for the pandemic clone, *E. coli* ST131. Further compounding this problem is the fact that many studies are conducted using common strains of pathogenic or avirluent bacteria such as CFT073, UTI89, or MG1655 a strain of K-12 (respectively), and not enough consideration has been given to clinical isolates of bacteria which each possess their own unique characteristics [21–23].

In one study evaluating CTX-M-14 and CTX-M-15 β-lactamase production amongst clinical isolates of *E. coli* it as determined that XQ12 and RS059 produced relatively the same amounts of CTX-M-15, despite XQ12 inducing pyelonephritis in 90% of animals tested in our study while RS059 only showed 50% infectivity [20]. CUMC247, CMB106, and JJ2244 also showed similar production of CTX-M-15 even though CUMC247 did not colonize any bladder tissues, while CMB106 and JJ2244 were proficient colonizers infecting 70% and 80% of kidneys respectively [20]. RS135 produced the highest level of CTX-M-15 in the study by Geyer et al., (2016) while being unable to cause either kidney or bladder infection in mice. Regarding CTX-M-14 β-lactamase production, C14 and Lo14 produced equivalent amounts of enzyme, while C14 was a prodigious infector of bladder and kidney tissues, while Lo14 was detectable in only one kidney and no bladder tissues in our study [20]. Taken together, although production of CTX-M-14 and CTX-M-15 have been considered virulence factors in ExPEC isolates, especially ST131 *E. coli*, their production alone does not predict their ability to cause pyelonephritis in a mouse model of transcending UTI.

Looking independently at phenotypic assays, clinical isolates showed less motility and biofilm formation than nonpathogenic lab strains. Of the clinical isolates, ST131 strains showed more motility and produced more biofilm than the non-ST131 clinical isolates. These results agree with the findings of Kondratyeva et al., (2017) who determined seven pathogenic strains of ST131 *E. coli* were more motile than 7 comparable strains of Non-ST131 MDR ESBL-producing clinical ExPEC *E. coli* isolates [24]. Given the increasing pandemic incidence of UTIs caused by ST131 isolates it was not surprising that these isolates would demonstrate greater motility and biofilm forming capabilities than their non-ST131 counterparts with motility and biofilm formation having been identified as mechanisms of pathogenesis [25]. However, a similar study comparing UTI clinical isolates of several sequence types, found multiple isolates were non-motile after 16 hours at 30°C with an equal number hypermobile as compared to CFT073; whereas, our study found most clinical isolates more motile than CFT073, with only 3/20 isolates less motile than the lab strain BW25113 or pathogenic CFT073 [14]. The same study found most clinical isolates tested produced less biofilm than CFT073 at 37°C; whereas, in our study we found CFT073 to be comparable to the ST131 clinical isolates yet lower than the lab strains for both static and shaking conditions [14]. The only ST131 isolate tested (HM6) also showed comparable biofilm production to CFT073 in the previous study [14]. These differences could be due to the divergence of experimental parameters with different temperatures and assay times but are also reflective of the variable nature of clinical isolates in phenotypic testing as noted throughout this paper. An unexpected finding from this study was the differential effect of static vs mid-log growth conditions for the outcomes of these assays. Static conditions are considered crucial for the production of type 1 fimbriae and the ability of pathogenic *E. coli* to produce biofilm, hemagglutinate, adhere to, and invade host tissues [17]. However, in our experiments, biofilm production increased under shaking conditions. These findings could point to other adhesins being important in biofilm production including other classes of fimbriae and pili, along with adherence-associated surface proteins such as OmpA to establish and maintain infections in ST131 *E. coli* [26].

Most *in vitro* assays for the evaluation of attachment to epithelial cells are done using static cultures [27]. Static cultures were first used to evaluate fimbriae because static cultures demonstrated more fimbriae production compared to shaking cultures making it easier to evaluate the protein structure [28]. We examined attachment of *E. coli* ST131 by evaluating fimbriae, curli, and p-pilus using *in vitro* assays with cultures grown in both static and shaking conditions. We reasoned that in a urinary tract infection it would be unlikely that the *E. coli* would exhibit a static phenotype. Overall ST131 clinical isolates were better able to adhere to, and invade, T24 human epithelial cell cultures than non-ST131 isolates regardless of static or shaking conditions. Although non-ST131 isolates collected from human urine showed a decrease ability to adhere to and invade epithelial cells they were still capable, while non-pathogenic ST10 lab strains showed limited to no ability for either. These results are corroborated by other phenotypic studies wherein the ST131 isolate had one of the highest association percentages of the clinical isolates tested; however, in that study the lab strain K12 also showed high cell association whereas our K12-derived lab strains did not [14]. The ability of non-ST131 isolates to adhere and colonize, albeit less successfully than ST131 isolates, suggests pathogenic mechanisms may have evolved in ST131 isolates enabling them to establish a strong pandemic prevalence [29]. In our experiments, we noted an increase in epithelial cell invasion under static conditions, but epithelial cell adhesion was variable depending on the isolate, many showing increased adhesion when grown to mid-log prior to co-culture.

Interestingly in our study, ST131 isolates were largely unable to cause hemagglutination in erythrocytes, commonly thought to be directly linked to pathogenicity through the expression of type 1 fimbriae in colonizing host tissues [17]. The only ST131 isolate that correlated with pathogenicity and infectivity in the mouse model and hemagglutination assay was isolate W15. But this correlation was only observed when cells were grown statically, not when grown to mid-log under shaking conditions. In contrast, ST10 lab strains, CFT073, and non-ST131 clinical isolates were capable of hemagglutination under static conditions [17]. CFT073 was also capable of hemagglutination using mid-log cells from a shaking culture. Similarly, in another study of clinical isolates, some isolates were capable of causing hemagglutination, while others were not; however, in that study the only ST131 evaluated showed strong hemagglutination capabilities whereas ours overwhelmingly did not [14]. These findings were unexpected and indicated the importance of other adhesins in the pathogenic phenotype of *E. coli* ST131 such as p-fimbriae and curli [17].

ST131 *E. coli* are often discussed as a group with similar traits, such as antibiotic resistance and some virulence factors. However, WGS of ST131 isolates has confirmed that these strains also have differences with respect to both resistance mechanisms and virulence factors. Johnson and colleagues have identified a common set of virulence factors (*fimH, fyuA, malX, usp, ompT, papA, kpsM* and *iutA*) that are often found in ST131 [30]. However, not all ST131 isolates have this same common set. Many of the genes encoding these virulence factors are found on the chromosome and constitute part of the core genome of *E. coli*; however, additional copies of these gene operons can be found in some isolates due to horizontal gene transfer and some lack the specific genes due to deletion events [31]. With the average *E. coli* genome consisting of between 4,000-5,000 genes, ST131 *E. coli* share a core genome of 3,712 genes with over 22,000 accessory genes identified across isolates. [31,32]. With the extreme variability in ST131 *E. coli* genomes, researchers turn to *in vitro* assays to help evaluate fitness and virulence of these isolates.

The overall objective of this research was to determine the correlation of commonly used *in vitro* phenotypic assays as predictive indicators for bacterial pathogenicity *in vivo*. At first glance, it appeared that hemagglutination and biofilm production had a negative association to pathogenicity, whereas motility, adhesion, and invasion positively correlated with infectivity. However, the statistical analyses of these findings differed. With Pearson correlations confirming the strongest association at a coefficient of 0.21, it is impossible to say that the *in vitro* results are definitively correlated to or predictive of *in vivo* outcomes. Our results are supported by another recent study where investigators determined weak Pearson correlation values for *in vitro* phenotypic assays to predict *in vivo* bladder colonization in a mouse model [14]. Between our data set and that of Shea, we had a slightly higher correlation for both motility and urine colonization (0.1349 vs 0.08827) as well as hemagglutination and urine colonization (0.2067 vs 0.1864). This could be due in part that our experiments focused primarily on ST131 *E. coli*, while Shea et al., (2022) screened a different subset of UPEC isolates including only one ST131 and 12 non-ST131s, or because we allowed our mouse model of UTI to develop for 28 days while Shea et al., (2022) ended their study at 48 hours post-inoculation. Therefore our mechanisms could point more toward the bacterial requirements necessary to maintain an infection while the other results indicate the necessary mechanisms to establish an infection [14]. Taken together, these two studies show that regardless of the *E. coli* isolates being evaluated, *in vitro* assays may be useful in determining phenotypic differences between isolates of *E. coli* but their usefulness in predicting pathogenicity is currently limited.

Although the literature tends to evaluate ST131 *E. coli* as a single pandemic clone it is clear from this study as well as WGS data that these organisms differ in their genomic makeup and their infectivity in a mouse UTI model. Expression of both CTX-M β-lactamases and OmpC porin production have also been shown to vary among strains of ST131, suggesting the physiology of these organisms differ [20,33]. Together, these data identify the uniqueness of individual clinical isolates of ST131. Further analyses on this group of *E. coli* are required with emphasis on physiological differences and how those differences impact pathogenesis. Differences that are uncovered may be key in determining how members of this clonal group have become such a dominant global pathogen.

## Materials and methods

### Ethics statement

All procedures involving animals were approved and in compliance with the Guide for the Care and Use of Laboratory Animals (protocol numbers 1174) by the Creighton University Institutional Animal Care and Use Committee.

### Cell and bacterial culture conditions

T24 human epithelial bladder cells (ATCC HTB-4) were grown in McCoy’s 5A modified medium (Gibco 16-600-082) with 10% fetal bovine serum (FBS) and 1% penicillin/streptomycin antibiotics at 37°C in the presence of 5% CO_2_. All bacteria strains used in this study are listed in Table 1. *E. coli* isolates were grown in Mueller-Hinton Broth (MHB) for growth curves, or Luria Broth (LB) for phenotypic analysis. LB contains 10g of tryptone and sodium chloride and 5g of yeast extract per liter. MHB is commonly used for antimicrobial susceptibility testing and conforms to the National Committee for Clinical Laboratory Standards recommended guidelines for cation composition (Ca2+ 20-25mg/L and Mg2+ 10-12.5mg/L).

### Murine model of urinary tract infection

Female C57BL/6 mice aged 42-48 were purchased from Charles River Laboratories for this study. A minimum of 6 mice were used for each bacterial strain. Animals were co-housed and provided food and water ad libitum. To ensure the urinary CFUs measured were due to the bacteria inoculated, isolates were subjected to phenotypic analysis (growth on MacConkey selective differential media and MIC measurements with Ampicillin and Cefotaxime) and CFUs determined based on the intended inoculum. To initiate ascending urinary tract infections mice were anesthetized with isoflurane and inoculated via transurethral catheterization as previously described [34]. Briefly, Catheters were made by attaching polyethylene tubing (BD Intramedic #22-204008 I.D. 0.011 mm x O.D. 0.28 mm) to a Sub-Q needle (Becton Dickinson #305115 26G 5/8”) on a 1 mL Leur-lock syringe (Fisherbrand #14955464). Bacteria were grown to mid-log phase in MHB shaking at 37 °C. Cultured bacteria were diluted to a 0.5 McFarland in sterile saline to a density of 1×10^8^ *E. coli* suspended in 50µL of saline as the inoculum. Prior to catheterization, urine was collected and plated on blood agar to detect the presence of any preexisting urine contamination. Urine was collected on Days 1, 3, 7, 10, 14, 21, and 28 post inoculation. Collected urine was serially diluted and 10 µL plated on LB agar in duplicate to count colony forming units (CFUs). On day 28, mice were euthanized and aseptically dissected. Blood was collected via cardiac puncture with an insulin syringe, photographs of kidneys were taken *in situ*, then bladder and kidneys were removed. Half of each kidney was immediately placed into a 1.5 mL tube and stored at −80°C for future use. The other half of each kidney and the bladder were placed into individual tubes containing 500 µL PBS and homogenized using zirconia/silica beads (2.3 mm diameter, BioSpec #11079125z). Homogenized tissues were serially diluted and 10 µL plated on LB agar square grid plates in duplicate to determine CFUs. Remaining tissue homogenate was stored at −80°C for future experiments.

### Bacteria strain phenotyping

#### Phenotypic Photos and Growth Curves

Bacteria strains were grown on blood agar plates overnight at 37°C. Colonies were taken from the blood agar plates and inoculated into 95 mL MHB sidearm flasks to a 0.1 OD600. Cultures were incubated at 37°C, shaking, with OD measurements taken at 15-minute intervals for 165 minutes. At 0.5 OD600, samples were taken and serially diluted to determine mid-log CFUs.

#### Motility

Assays were performed as previously described with modifications [14]. Briefly, overnight cultures of each strain grown on blood agar plates were diluted to a 2.0 McFarland in sterile saline and 10 µL of the suspension was inoculated directly into the center of a motility plate (0.3% bactoagar w/v in LB). Bacteria were cultured for 10 hours at 37°C, photographs taken and the diameter of the zone of growth was measured in mm.

#### Biofilm Formation

Biofilm was measure for each isolate in static and shaking conditions as previously described [35]. Briefly, bacteria strains were cultured in overnight static conditions or shaking to mid-log phase at 37°C in LB. Cultures were diluted to 0.5 McFarland and 100 µL of suspension was added to a 96-well plate containing 100 µL of LB. Plates were incubated at 37°C either statically or shaking, respectively, for 24 hours. The media was removed, and wells were washed with sterile water to remove non-adherent bacteria. Plates were stained with 200 µL Crystal Violet for 15 minutes. Crystal violet was removed, and plates were washed three times with sterile water and left upside down to dry overnight. After drying, stained biofilm was solubilized with 200 µL 30% acetic acid for fifteen minutes. Absorbance was read at 575 nm and blank-adjusted. Biofilm production was compared to the absorbance of a well incubated with only LB and classified as strong (OD>2×ODc), moderate (1.5×ODc<OD≤2×ODc), weak (ODc<OD≤1.5×ODc), or absent (OD≤ODc) (Zhang et al., 2021).

#### Adhesion and invasion

Human bladder epithelium T24 cells (ATCC #HTB-4) were grown to confluence in 24-well plates. Cells were trypsinized and counted using trypan blue and a Countess 3 (ThermoFisher #A50298). Using cell counts, a multiplicity of infection of 100 was determined. Bacteria were grown statically or shaking to mid-log in LB and diluted to a 0.5 McFarland in sterile saline. The appropriate volume of suspension was pelleted by centrifugation and resuspended in 1 mL of serum-free, antibiotic-free, media. T24 cell culture media was removed, cells were washed with PBS, and bacterial suspension media was added and incubated for 2 hours at 37°C 5% carbon dioxide. During this time, bacterial inoculum was serially diluted and plated to determine inoculum CFUs. After incubation, media was removed, and cells were washed three times with PBS to remove any non-adherent bacteria. Epithelial cells were lysed with 0.4% Triton-X 100 in PBS for 30 minutes at 4°C shaking. The suspension was serially diluted and plated on LB agar for CFUs. Adherent bacteria are reported as percent CFUs of inoculum. For the invasion assay, T24 cells were cultured with static or shaking bacteria at a MOI 100 as described above. Prior to epithelia cell lysis, T24 cells and adherent bacteria were cultured with media containing Polymyxin B (100µg/mL) for one and a half hours to kill extracellular bacteria [36]. Cultures were then washed with PBS, lysed, and plated as described above. Intracellular bacteria are reported as percent CFUs of inoculum.

#### Hemagglutination

Overnight cultures were pelleted and resuspended in PBS, then serially diluted (no dilution, 1:1 to 1:64) and plated in 96-well round bottom culture plates. Samples were co-cultured with 3% guinea pig erythrocytes. Mannose was added to the erythrocytes and undiluted culture in the last row of the plate to act as a competitive binding protein for type 1 fimbriae. Bacteria were cultured either statically for 24 hours or shaking to mid-log phase. Mannose sensitive hemagglutination was evaluated as previously described [37]. Briefly, cultures were diluted to 0.5 OD600, centrifuged to pellet, and resuspended in 75 µL PBS. Bacteria suspension was added to the first row in a 96-well plate and serially diluted 1:2 in following rows. Guinea pig erythrocytes were washed in PBS and resuspended at 3% vol/vol in PBS. 25 µL red blood cell suspension was added to each row of 50 µL bacterial suspensions, in the final row mannose was added to a final concentration of 50mM to inhibit type 1 fimbriae binding. Plates were incubated at 37°C shaking for 5 minutes then transferred to 4°C for one hour. Titer was determined as the last dilution where hemagglutination was observed.

#### Curli production

Congo Red Assay Plates were made as previously described [18]. In brief, plates were made with 15g/L Agarose, 15.5g/L Low-Salt LB Broth, and a final concentration of 20mg/L Coomassie Brilliant Blue and 40mg/L Congo Red. Bacteria were grown overnight in LB broth under either shaking or static conditions at 37°C. 3µL of each isolate were spot plated in triplicate onto two different Congo Red plates. One plate was incubated at 37°C for 24 hours, while the other was incubated for 48 hours at 27°C. Colony color and morphology were scored as previously described by three independent evaluators and then aggregated. Colonies were scored as follows: Score 0 = an all-white colony, Score 1 = light color, Score 2 = some dark color or ring structures with a matte surface just where the color is, Score 3 = a dark, even color accompanied by a matte/dry surface, Score 4 = dark color and a rough, wrinkled colony [18].

### Statistical analysis

Data are presented as mean value of three replicates. Statistical analyses were conducted in GraphPad Prism version 9.5.1. For phenotypic assays and bacterial load, one-way analysis of variance (ANOVA) with post-hoc Tukey tests were conducted to compare differences between strains. For mouse studies, proportions were compared using two-tailed Fisher’s exact tests. Pearson correlations were conducted between all phenotypic and *in vivo* datasets individually and compared using multiple linear regressions. The threshold for statistical significance for all analyses was set at *p*≤0.05.

## Acknowledgements

We thank Kori Klingelsmith and Stacey Morrow for excellent technical assistance. We thank Dr Heike Brötz-Oesterhelt for the OmpC/OmpF knockout strain, Department of Microbial Bioactive Compounds, Interfaculty Institute of Microbiology and Infection Medicine, University of Tübingen, D-72076 Tübingen, Germany; German Center for Infection Research (DZIF) Partner Site, D-72076 Tübingen, Germany.

We thank the Creighton University Animal Research Facility for excellent animal care.

## Author Contributions

**Conceptualization:** Courtney P. Rudick, Rachel S. Cox, Travis J. Bourret, Nancy D. Hanson.

**Data curation:** Courtney P. Rudick, Rachel S. Cox.

**Formal analysis:** Courtney P. Rudick.

**Funding acquisition:** Nancy D. Hanson.

**Investigation:** Courtney P. Rudick, Rachel S. Cox.

**Methodology:** Courtney P. Rudick, Rachel S. Cox, Travis J. Bourret.

**Project administration:** Nancy D. Hanson.

**Supervision:** Travis J. Bourret, Nancy D. Hanson.

**Validation:** Courtney P. Rudick, Travis J. Bourret, Nancy D. Hanson.

**Writing – original draft:** Courtney P. Rudick

**Writing – review & editing:** Courtney P. Rudick, Rachel S. Cox, Travis J. Bourret, Nancy D. Hanson.

## Supporting information

**S1 Fig. Growth curves for all *E. coli* isolates used in this study.** Isolate growth curves show no differences that would account for the differences in phenotype assay results or mouse model infectivity. O.D.: Optical Density at 600nm.

**S2 Fig. Urine colonization throughout study.** Urine was collected to determine colony forming units. (A) All isolates (B) BWΔ*ompF* (C) W15 (D) C15 (E) C14. Graph line and point color represent isolate sequence type as noted in the figure legend.

**S3 Fig. Motility assay plate photographs.** Motility assays were performed by inoculating soft agar plates (0.3% bactoagar w/v in LB) with a 10µL solution of overnight cultures of each strain which were grown on blood agar plates and diluted to a 2.0 McFarland in sterile saline. Plates were incubated for 10 hours at 37°C and then zone of growth diameter was measured in mm. Measurements for each isolate are listed in Figure 4.

**S4 Fig. Linear regressions for phenotypic assays and *in vivo* infection results.** Linear regressions of motility assays compared to (A) urine CFUs. (B) bladder CFUs, and (C) kidney CFUs. Linear regressions of hemagglutination assays compared to (D) Urine CFUs. (E) Bladder CFUs, and (F) Kidney CFUs. Linear regressions of biofilm production compared to (G) Urine CFUs. (H) Bladder CFUs, and (I) Kidney CFUs. Linear regressions of T24 epithelial cell adhesion assays compared to (J) Urine CFUs. (K) Bladder CFUs, and (L) Kidney CFUs. Linear regressions of invasion assays compared to (M) Urine CFUs. (N) Bladder CFUs, and (O) Kidney CFUs. Linear regressions of congo red curli assays compared to (P) Urine CFUs. (Q) Bladder CFUs, and (R) Kidney CFUs.

